# Quantifying the impact of genotype-dependent gene flow on mutation fixation in subdivided populations

**DOI:** 10.1101/2023.11.29.569213

**Authors:** Loïc Marrec

## Abstract

In the wild, any population is likely to be spatially structured. Whereas we deeply understand evolutionary dynamics in well-mixed populations, our understanding of evolutionary dynamics in subdivided populations needs to be improved. In this work, I quantify the impact of genotype-dependent gene flow on the evolutionary dynamics of a subdivided population. Specifically, I build a model of a population structured as the island or the stepping stone model in which genotype-dependent gene flow is represented by individuals migrating between its sub-populations at a rate depending on their genotype. I analytically calculate the fixation probability and time of a mutation arising in the subdivided population under the low migration limit, which I validate with numerical simulations. I find that the island and the stepping stone models lead to the same fixation probability. Moreover, comparing the fixation probability in these models to the one in a well-mixed population of the same total census size allows me to identify an effective selection coefficient and population size. In the island and the stepping stone models, the effective selection coefficient differs from the selection coefficient if the wild-type and the mutant migration rates are different, whereas the effective population size equals the total census size. Finally, I show that genotype-dependent gene flow increases the fixation time, which allows for distinguishing the island and the stepping stone models, as opposed to the fixation probability.

## Introduction

In the wild, any population has a certain degree of spatial structure. For example, human activity leads to fragmentations of natural habitats, causing animal populations to be subdivided into sub-populations [1, 2]. Although habitats resulting from fragmentations are isolated, some individuals may migrate from one sub-population to another, a process called gene flow [3]. Like mutations, gene flow is an important mechanism in evolutionary biology as it increases genetic diversity, as opposed to natural selection and genetic drift. However, gene flow theoretically has a disruptive effect on adaptation by counteracting selection and promoting the fixation of deleterious mutations [4], although some empirical evidence showed the opposite [5]. Thus, whereas we understand the impact of natural selection and genetic drift on evolutionary dynamics in a well-mixed population (at least in the absence of epistasis [6]), considering spatial structure and gene flow makes predicting evolutionary outcomes more challenging.

This challenge has been addressed since the beginning of theoretical population genetics by calculating the fixation probability of a mutation in a subdivided population [7]. It was long believed that most spatial structures would give the same fixation probability as in well-mixed populations [8, 9, 10, 11, 12]. However, further studies pointed out that this result emerges from the conservative gene flow assumption (i.e., migration of individuals does not change sub-population sizes), and challenged this long-standing belief by considering more complex cases. For example, a meta-population model, including local extinctions and colonizations, led to a fixation probability of beneficial mutations different from that in well-mixed populations [13]. A more recent model showed that some spatial structures could either decrease the fixation probability of deleterious mutations and increase that of beneficial mutations or do the opposite depending on the gene flow pattern [14], which indicates that some spatial structures impact the efficacy of natural selection.

The efficacy of natural selection in subdivided populations is related to the effective population size [15, 16]. As explained in [17], if the effective population size of a spatially structured population is larger than its total census size, the structure increases the efficacy of natural selection, i.e., it increases the fixation probability of beneficial mutations and decreases that of deleterious mutations compared to a well-mixed population. Conversely, if the effective population size of a spatially structured population is lower than its total census size, the structure decreases the efficacy of natural selection, i.e., it decreases the fixation probability of beneficial mutations and increases that of deleterious mutations compared to a well-mixed population. This link between the effective population size and the fixation probability of a mutation in a subdivided population explains why so much effort has gone into deriving the former [17, 18]. Whereas the impact of the population structure topology on evolutionary dynamics has received much attention, for example, through evolutionary graph theory [20], much less is known about the impact of genotype-dependent gene flow. In particular, numerous theoretical studies investigating evolutionary dynamics in subdivided populations assume genotype-independent gene flow [3, 4, 21]. Yet, there is empirical evidence that genotype-dependent gene flow occurs in nature [22], e.g., in aquatic species [23, 24], butterflies [25, 26], and plants [27]. For example, Glanville fritillary butterflies (*Melitaea cinxia*) carrying a specific allele of the metabolic enzyme phosphoglucose isomerase have higher metabolic flight rates and, thus, higher dispersal rates [25]. Therefore, there is a need to better understand the impact of genotype-dependent gene flow on the evolutionary dynamics of subdivided populations.

In this paper, I investigate the evolutionary dynamics of a subdivided population, focusing on the impact of genotype-dependent gene flow on their evolutionary outcome. To do so, I build a model describing a population subdivided as the island or the stepping stone model in which gene flow is modeled by individuals migrating between the sub-populations at a rate depending on their genotype. To quantify the evolutionary dynamics of the subdivided population, I derive analytical predictions for the fixation probability, the number of (fixed) migrants, and the fixation time of a mutation, which I compare to numerical simulations.

## Model and Methods

### Subdivided population model

I build a continuous-time model with overlapping generations in which each life cycle event (i.e., reproduction, death, migration) is decoupled from each other and occurs at random. I consider an asexual haploid population of total census size *N*_tot_ subdivided into *D* well-mixed (or homogeneous) demes whose census sizes, although varying over time, are limited by a carrying capacity *N*. The carrying capacity can result from, for example, limited space or nutrients. I focus on a single locus where two alleles exist, namely wild-type (W) and mutant (M), resulting in genotypes whose intrinsic birth rates are denoted by *b*_W_ = 1 and *b*_M_ = 1 + *s*, respectively, where *s* is the selection coefficient. The sign of the selection coefficient *s* describes whether the mutation is beneficial (i.e., *s >* 0), neutral (i.e., *s* = 0), or deleterious (i.e., *s <* 0). I also define an intrinsic death rate *d*, identical for both genotypes, and genotype-dependent migration rates per individual, denoted by *m*_W_ and *m*_M_. I do not consider *de novo* mutations, which is equivalent to considering a zero mutation probability upon reproduction. Similarly to [11, 14, 28, 29], I assume a low migration limit so that no migration occurs during the fixation of either genotype in a deme. In this limit, the migration rate is much lower than the fixation rate, so each deme can be assumed to be fully wild-type or mutant most of the time. Within each deme, the population follows a continuous-time logistic growth with density-dependent birth rates and density-independent death rates. More specifically, the wild-type and mutant *per capita* birth rates satisfy *b*_W_(1 − *N*_W_*/N*) and *b*_M_(1 − *N*_M_*/N*), respectively, whereas the wild-type and mutant *per capita* death rates are equal to *d*. I focus on the saturation phase in which the census size fluctuates around its equilibrium, that is, *N*_W_ = *N* (1 − *d/b*_W_) for the wild-type demes and *N*_M_ = *N* (1 − *d/b*_M_) for the mutant demes. I consider timescales much shorter than those of extinctions resulting from demographic stochasticity [30].

### Fixation dynamics in a single deme

I shall recall some known results about the fixation dynamics of either genotype in a single well-mixed deme assuming no migration, which will be useful for the following analytical derivations. Although I deal with size-varying demes, their size fluctuates around their equilibrium, which allows me to use the Moran process to derive the fixation probabilities and times [31]. Suppose I consider a wild-type deme of size *N*_W_ in which there is a single mutant. This mutant takes over the deme with a probability equal to

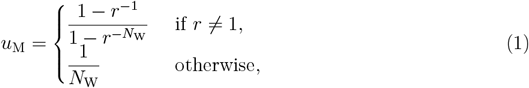

where *r* = 1 + *s* is the relative fitness of the mutant (see the supplement of [32] for a full derivation). Similarly, the fixation probability of a wild type in a mutant deme of size *N*_M_ is given by

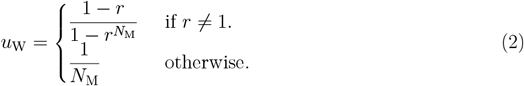

Applying the weak selection assumption to the fixation probabilities *u*_W_ and *u*_M_ (i.e., |*s*| ≫ 1), in addition to considering that the deme sizes satisfy 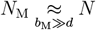 and 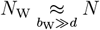, leads to

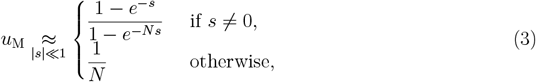

and

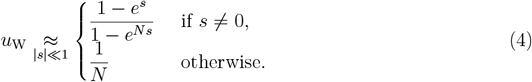

The form of Equations 3 and 4 is similar to the well-known fixation probability in a Wright-Fisher population obtained in the Kimura’s diffusion limit [33, 31]. Both fixation probabilities can be compared by computing the ratio *u*_W_*/u*_M_, which assuming *N* |*s*| *≫* 1 gives

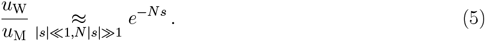

Since I want to focus on the low migration limit so that no migration disturbs a fixation process within a deme, I need to recall the mean fixation time of either genotype, namely wildtype and mutant. The fixation time of a mutant in a wild-type deme reads (see the supplement of [32] for a full derivation)

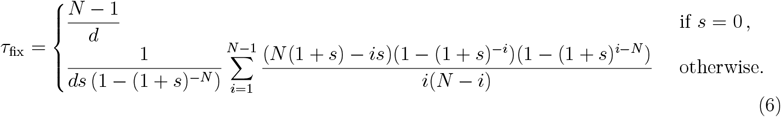

Similarly, one can compute the mean fixation time of a wild type in a mutant deme, but it is equal to that of a mutant in a wild-type deme. In other words, the fixation time of a mutation is the same as for a wild type with the same strength of selection (i.e., |*s*|), a result shown in [9].

### Fixation probability in the well-mixed model

Before considering spatially structured populations, I shall recall the fixation probability of a mutant fraction *p* in a well-mixed population of total census size *N*_tot_ (see Figure 1 left), which reads

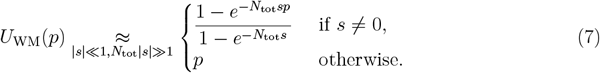

The fixation probability of a mutation in a well-mixed population will serve as a basis to assess the impact of spatial structure on the evolutionary dynamics of subdivided populations.

**Figure 1:**
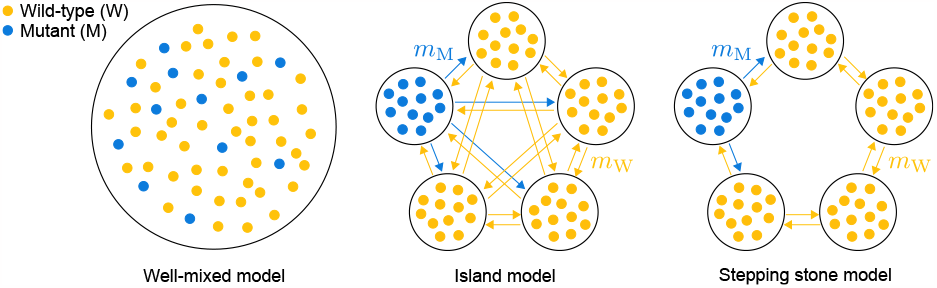
Some examples of subdivided populations. The well-mixed population model and two models of populations that are subdivided into a finite number of demes in which there are two genotypes, namely wild-type (W) and mutant (M). The island model represents a subdivided population in which each deme is connected to all the others, allowing its individuals to migrate between any pair of demes at a rate depending on their genotype (i.e., *m*_M_ and *m*_W_ for the mutants and the wild types, respectively). The stepping stone model represents a subdivided population arranged on a ring in which each deme is connected to its two adjacent neighbors, allowing its individuals to migrate between adjacent demes at a rate depending on their genotype (i.e., *m*_M_ and *m*_W_ for the mutants and the wild types, respectively). Parameter values: number of demes *D* = 5, wild-type deme size *N*_W_ = 12, mutant deme size *N*_M_ = 12, total census size *N*_tot_ = 60.

### Fixation dynamics in the island model

The island model is a subdivided population model in which each deme is connected to all the others, resulting in *D*(*D* − 1) migration paths (see Figure 1 center). In the island model, the fixation probability of a mutation starting from a fraction of fully mutant demes *p* reads in the low migration limit

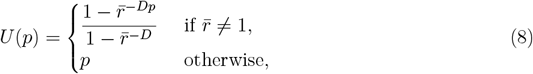

where 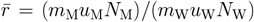 is the relative fitness of mutant demes (Equation 8 was derived in [14] but I extended it to genotype-dependent migration rates). I can simplify Equation 8 by considering that *b*_M_ ≫ *d* and *b*_W_ ≫ *d*, which implies *N*_M_ ≈ *N* and *N*_W_ ≈ *N*, as well as |*s*| 1 and *N* |*s*| ≫1, which implies *u*_W_*/u*_M_ ≈*e*^*−Ns*^. These assumptions allow me to write Equation 8 as

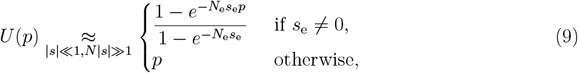

where *N*_e_ = *D × N* = *N*_tot_ is the effective population size and *s*_e_ = *s* − log(*α*)*/N* is the effective selection coefficient with *α* = *m*_W_*/m*_M_. I defined the effective population size and selection coefficient, i.e., *N*_e_ and *s*_e_, respectively, to make the fixation probability *U* have the same form as *U*_WM_ (see Equations 7 and 9).

To further characterize the dynamics of mutation fixation, I want to compute the mean number of fixed migrants, regardless of their genotype, until the fixation of a mutation, given this mutation becomes fixed. More specifically, a fixed migrant is a migrant fixing in a deme of the other genotype, leading to a change in the number of wild-type and mutant demes. To compute the number of fixed migrants, I analyze the evolutionary dynamics of the subdivided population by using a Markov process tracking the number of mutants demes *i* [34, 35]. Since I focus on the low migration limit, only two events can change the number of mutant demes. Either a wild type migrates from one of the *D* − *i* wild-type demes to one of the *i* mutant demes and becomes fixed, decreasing the number of mutant demes by 1 (i.e., *i* → *i* − 1), or a mutant migrates from one the *i* mutant demes to one of the *D* − *i* wild-type demes and becomes fixed, increasing the number of mutant demes by 1 (i.e., *i* → *i* + 1). The probabilities that these events occur upon a migration leading to a fixation within a deme are given by

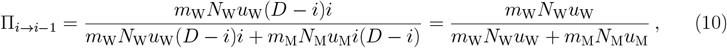

and

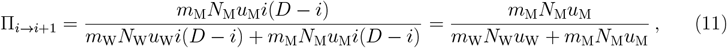

respectively. Then, using Equation 1.39 from [36] allows me to compute the mean number of fixed migrants *n*_fix_ starting from a fraction of fully mutant demes *p* until the fixation of a mutation, given it fixes, which reads

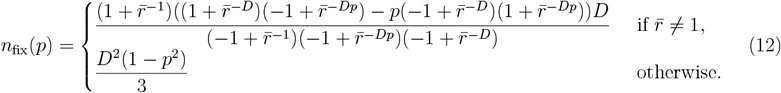

An additional interesting quantity to characterize the dynamics of mutation fixation is the mean number of migrants, which, as opposed to the mean number of fixed migrants, does not distinguish between migrants fixing and those going extinct (i.e., does not distinguish between migrants leading to a change in the number of wild-type and mutant demes and those leaving the number of wild-type and mutant demes unchanged). In this case, I need to modify the probabilities I introduced earlier. Although the events changing the number of mutant demes are still the same, the probabilities that these events occur upon a migration, regardless of whether the migrant fixes or goes extinct, are given by

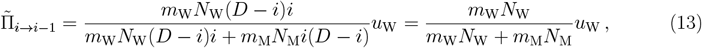

and

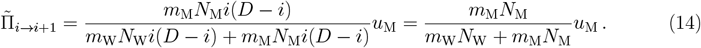

By applying the same method as for the mean number of fixed migrants (i.e., by using Equation 1.39 of [36]), I obtain the mean number of migrants until the fixation of a mutation, given it fixes, which reads

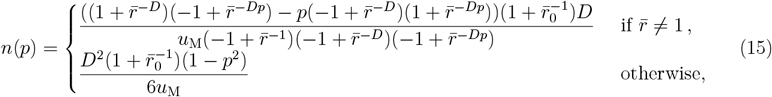

where 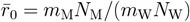.

To quantify the timescales associated with mutation fixation, I compute the mean fixation time of a mutation, given it fixes. The reasoning is similar to the one I applied for the number of (fixed) migrants, but here, I employ the transition rates rather than the probabilities. The transition rates satisfy

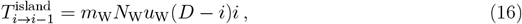

and

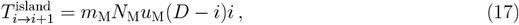

and lead to the fixation time

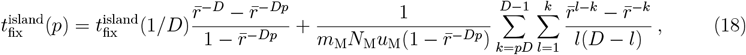

where

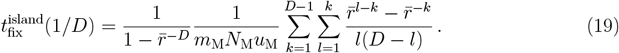

Finally, to make sure I choose parameter values corresponding to the low migration limit, and also to rigorously compare different population structures, I need to compute the total migration rate in the island model. Since I consider genotype-dependent migration rates, the total migration rate will depend on the number of mutant demes in the subdivided population, which is why I focus on the total migration rate averaged over the number of mutant demes, which is given by

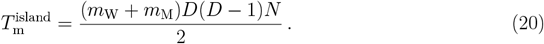

Note that I considered that *b*_M_ *≫ d* and *b*_W_ *≫ d*, which implies *N*_M_ *≈ N* and *N*_W_ *≈ N*.

### Fixation dynamics in the stepping stone model

The stepping stone model is a subdivided population model in which each deme is arranged on a ring and connected to their left and right neighbors, resulting in 2*D* migration paths (see Figure 1). The fixation probability in the stepping stone model starting from a fraction of fully mutant demes *p* in the low migration limit has exactly the same form as the island model (see Equation 8) [14].

Similarly, the probabilities that the number of mutant demes changes in the stepping stone model are the same as the island model (see Equations 10, 11, 13, and 14). This result may seem surprising since, in the island model, there are *i*(*D* − *i*) migration paths that can lead to a change in the number of mutant demes *i*, whereas there are 2 in the stepping stone model. However, in both models, the number of migration paths in the probability that the number of mutant demes increases by one is the same as the probability that the number of mutant demes decreases by 1, which makes it vanish. As a result, both models have the same mean number of fixed migrants *n*_fix_ and the same mean number of migrants *n*.

Now, I compute the mean fixation time of the mutation in the stepping stone model, given it fixes. The reasoning is similar to the one I applied for the island model, but the transition rates in the stepping stone model differ from those in the island model. The transition rates in the stepping stone model satisfy

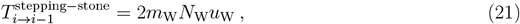

and

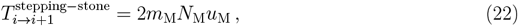

which lead to the fixation time

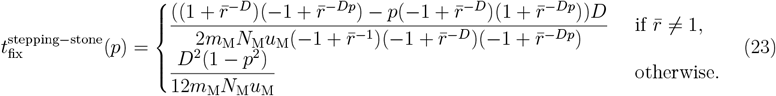

Similarly to the island model, I compute the total migration rate averaged over the number of mutant demes in the stepping stone model, which reads

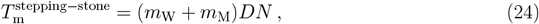

where I considered that *b*_M_ *≫ d* and *b*_W_ *≫ d*, which implies *N*_M_ *≈ N* and *N*_W_ *≈ N*.

### Numerical simulations

To ensure my analytical predictions are correct, I compare them to numerical simulations based on a Gillespie algorithm [37, 38]. Since I focus on the low migration limit, I can simply simulate the stochastic dynamics of the number of mutant demes, denoted by *i*. The elementary events that can happen are an increase or a decrease in the number of mutant demes (i.e., *i → i* + 1 and *i → i −* 1, respectively)

- 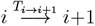: Increase in the number of mutant demes with rate 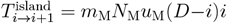 in the island model and rate 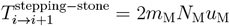 *m*_M_*N*_M_*u*_M_ in the stepping stone model.
- 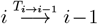: Decrease in the number of mutant demes with rate 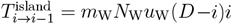 in the island model and 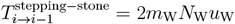 in the stepping stone model.

The total rate of events is given by *T*_*i*_ = *T*_*i→i*+1_ + *T*_*i→i−*1_, and simulation steps are as follows

1. Initialization: At time *t* = 0, I start with *D −* 1 wild-type demes and one mutant deme.
2. Monte Carlo step: Time *t* is incremented by Δ*t*, which is sampled from an exponential distribution with mean 1*/T*_*i*_. The next event to occur is chosen proportionally to its probability (i.e., *T*_*i*→*i*+1_*/T*_*i*_ and *T*_*i* →*i* 1_*/T*_*i*_ for an increase and a decrease in the number of mutant demes, respectively), and is executed.
3. I go back to Step 2 unless only one genotype, either wild-type or mutant, remains in the meta-population, which corresponds to the fixation of one genotype. In other words, simulation is ended when fixation occurs.

Simulations were performed with Matlab (version R2021a). Annotated codes to repeat the simulations and visualizations are available on GitHub (https://github.com/LcMrc). Throughout this work, I will consider parameter values such that the total migration rates averaged over the number of mutant demes in the island and the steeping stone models are equal (i.e., 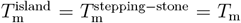; see Equations 20, and 24). Also, I will choose parameter values such that the fixation time of either genotype within a deme is much shorter than the time between two migrations (i.e., 1*/T*_m_ *≫ τ*_fix_; see Equations 6, 20, and 24).

## Results

### Fixation probability

The fixation probability is a key quantity in population genetics, as it determines whether a mutation will likely take over a population and, thus, reach a frequency of 1 [7]. My analytical predictions derived in the Model and Methods section allowed me to obtain an equation for the fixation probability *U* of a fraction *p* of mutants in a subdivided population of *D* demes of size *N*. The fixation probability *U* given by Equation 9 is the same for the island and the stepping stone models, and has the same form as the fixation probability in a Wright-Fisher population obtained in the Kimura’s diffusion limit [31, 33]. Therefore, the fixation probability is a quantity that does not allow for distinguishing between the two models. Interestingly, my analytical derivations enabled me to identify an effective selection coefficient *s*_e_ that differs from the selection coefficient *s* if the wild-type and mutant migration rates differ (i.e., *s*_e_ ≠ *s* if *m*_M_ ≠ *m*_W_). Thus, the well-known fixation probability of a neutral mutation, which is equal to the initial fraction of fully mutant demes *p*, is obtained when *s*_e_ = 0 rather than *s* = 0. As shown in Figure 2, genotype-dependent migration rates induce a shift in the fixation probability *U*, when plotted as a function of the selection coefficient *s*, toward lower or higher selection coefficient *s* depending on the value of the ratio *α* of the wild-type and mutant migration rates. This result highlights that the birth rates alone do not suffice to assess the fitness of a mutation in a subdivided population. For instance, a mutant reproducing less frequently than the wild type could still take over the entire population if it can migrate more often than the wild type.

**Figure 2:**
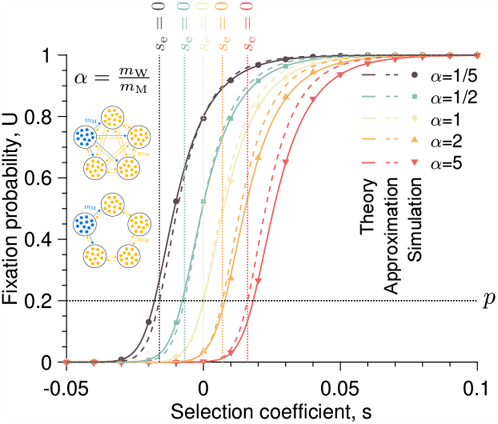
The fixation probability sheds light on an effective selection coefficient. **A** The fixation probability *U* of a mutation starting from one fully mutant deme is plotted against the selection coefficient *s* for different ratios *α* = *m*_W_*/m*_M_ of the wild-type and mutant migration rates. The solid lines represent the analytical predictions whereas the dashed lines show the approximated predictions (i.e., Equations 8 and 9, respectively). The markers are simulated data averaged over 10^4^ replicates. The vertical dotted lines show the effectively neutral case (i.e., *s*_e_ = 0, where *s*_e_ = *s* log(*α*)*/N* with *α* = *m*_W_*/m*_M_), whereas the horizontal dotted line corresponds to the fixation probability of a neutral mutation, that is, the initial fraction *p* of mutant demes. Parameter values: number of demes *D* = 5, deme size *N* = 100, wild-type birth rate *b*_W_ = 1, death rate *d* = 0.1, wild-type and mutant migration rates for the island model (*m*_W_, *m*_M_) × 10^*−*8^ = {(2.06, 10.3), (4.11, 8.23), (6.17, 6.17), (8.23, 4.11), (10.3, 2.06)} (from top to bottom), wild-type and mutant migration rates for the stepping stone model (*m*_W_, *m*_M_) × 10^*−*8^ = {(4.12, 20.6), (8.23, 16.5), (12.3, 12.3), (16.5, 8.23), (20.6, 4.12)} (from top to bottom).

Moreover, I find that the effective population size *N*_e_ equals the total census size *N*_tot_ = *D* × *N*, which shows that genotype-dependent migration rates do not decrease or increase the efficacy of natural selection in the island and the stepping stone models, as opposed to asymmetric migration rates in some subdivided populations [14]. The absence of decrease and increase of the efficacy of natural selection explains why the curves in Figure 2, which show the fixation probability *U* as a function of the selection coefficient *s* for different ratios *α* of the wild-type and mutant migration rates, have the same shape.

### Number of fixed migrants until mutation fixation

The fixation probability does not provide an exhaustive picture of the dynamics of mutation fixation. In particular, the fixation probability does not provide any insights into the timescales involved in evolutionary dynamics. Yet, the timescales associated with fixation processes are important to quantify. For example, comparing the timescale of mutation fixation to that of environmental changes allows for assessing the adaptation and persistence of a population undergoing changing environments. I found that the fixation probability is the same for the island and the stepping stone models, thus requiring additional quantities to distinguish them. For these reasons, I now focus on the number of fixed migrants, whether wild-type or mutant, until the fixation of a mutation, given this mutation gets fixed. As a reminder, a fixed migrant is a migrant fixing in a deme of the other genotype, leading to a change in the number of wild-type and mutant demes. As shown in Figure 3A and Equation 12, the number of fixed migrants *n*_fix_ is the same for the island and the stepping stone models, and also depends on an effective selection coefficient *s*_e_. Although the number of fixed migrants *n*_fix_ does not allow for distinguishing both structures, it provides insights into the mutation fixation dynamics. More specifically, the number of fixed migrants *n*_fix_ ranges from *D*(1 − *p*), obtained for large and low effective selection coefficients *s*_e_, to (*D*^2^ − 1)*/*3, obtained for the effectively neutral case (i.e., *s*_e_ = 0). This result confirms that the effectively neutral case is governed by strong stochasticity, resulting from genetic drift, and leads to several fixations of either genotype within demes before the mutation takes over the entire population, given the mutation becomes fixed. Conversely, the cases in which the effective selection coefficient *s*_e_ is nonzero are driven by the deterministic force of natural selection, which leads to a sequential fixation of mutants within each deme until the mutation takes over the entire population, while no wild type fixes.

**Figure 3:**
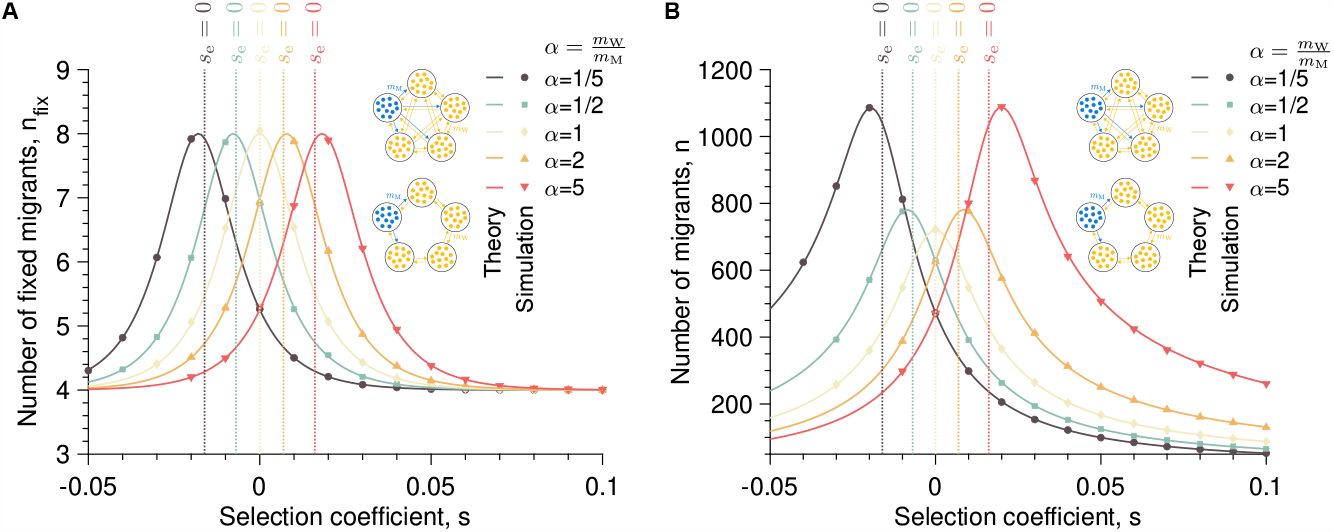
Genotype-dependent migration rates increase the number of migrants but not the number of fixed migrants. **A** The number of fixed migrants *n*_fix_ starting from one fully mutant deme is plotted against the selection coefficient *s* for different ratios *α* = *m*_W_*/m*_M_ of the wild-type and mutant migration rates. **B** The number of migrants *n* is plotted against the selection coefficient *s* for different ratios *α* = *m*_W_*/m*_M_ of the wild-type and mutant migration rates. In both panels, the solid lines represent the analytical predictions (i.e., Equations 12 and 15), the markers are simulated data averaged over 10^4^ replicates, and the vertical dotted lines show the effectively neutral case (i.e., *s*_e_ = 0, where *s*_e_ = *s* − log(*α*)*/N* with *α* = *m*_W_*/m*_M_). Parameter values: number of demes *D* = 5, deme size *N* = 100, wildtype birth rate *b*_W_ = 1, death rate *d* = 0.1, wild-type and mutant migration rates for the island model (*m*_W_, *m*_M_) × 10^*−*8^ = {(2.06, 10.3), (4.11, 8.23), (6.17, 6.17), (8.23, 4.11), (10.3, 2.06)} (from top to bottom), wild-type and mutant migration rates for the stepping stone model (*m*_W_, *m*_M_) × 10^*−*8^ = {(4.12, 20.6), (8.23, 16.5), (12.3, 12.3), (16.5, 8.23), (20.6, 4.12)} (from top to bottom).

### Number of migrants until mutation fixation

In the previous paragraph, I focused on the number of fixed migrants until the mutation takes over the entire population, given it becomes fixed. This quantity only provides a partial picture of migration events since it focuses only on migrations leading to a change in the number of mutant demes. Now, I turn my attention to the total number of migrants until the fixation of a mutation, given this mutation fixes, regardless of whether migrants lead to a change in the number of mutant demes. As shown in Figure 3 and Equation 15, the number of migrants *n* is the same for the island and the stepping stone models, as the fixation probability *U* and the number of fixed migrants *n*_fix_, and depends on an effective selection coefficient *s*_e_. However, as opposed to the number of fixed migrants *n*_fix_, the largest value of the number of migrants *n*, obtained for the effectively neutral case (i.e., *s*_e_ = 0), depends on the ratio *α* of the wild-type and mutant migration rates. More specifically, the larger the difference between the wild-type and mutant migration rates, i.e., *m*_W_ and *m*_M_, respectively, the larger the maximum value of the number of migrants *n* obtained for *s*_e_ = 0.

### Fixation time

Calculating the number of (fixed) migrants until the fixation of a mutation, given this mutation becomes fixed, allowed one to get insights into the evolutionary dynamics of subdivided populations. Now, I take a step further by considering the fixation time of a mutation. Figure 4 and Equations 18 and 23 show that the island and the stepping stone models have different fixation times, as opposed to the fixation probability and the number of (fixed) migrants. Thus, the fixation time is a quantity that allows for distinguishing both models. Specifically, as shown in Figure 4C, the fixation dynamics are faster in the island model than in the stepping stone model, although the ratio between both ranges only from 1.2 to 1.25. The difference between both models may appear surprising for two reasons: i) I set the same total migration rate for both models, and ii) I showed that both models have the same number of (fixed) migrants until mutation fixation (see Figure 3). This difference is due to the fact that the island and the stepping stone models do not have the same number of migration paths. For example, an island model with a single mutant deme has *D* − 1 migration paths to increase the number of mutant demes, whereas the stepping stone model has 2 migrations paths.

**Figure 4:**
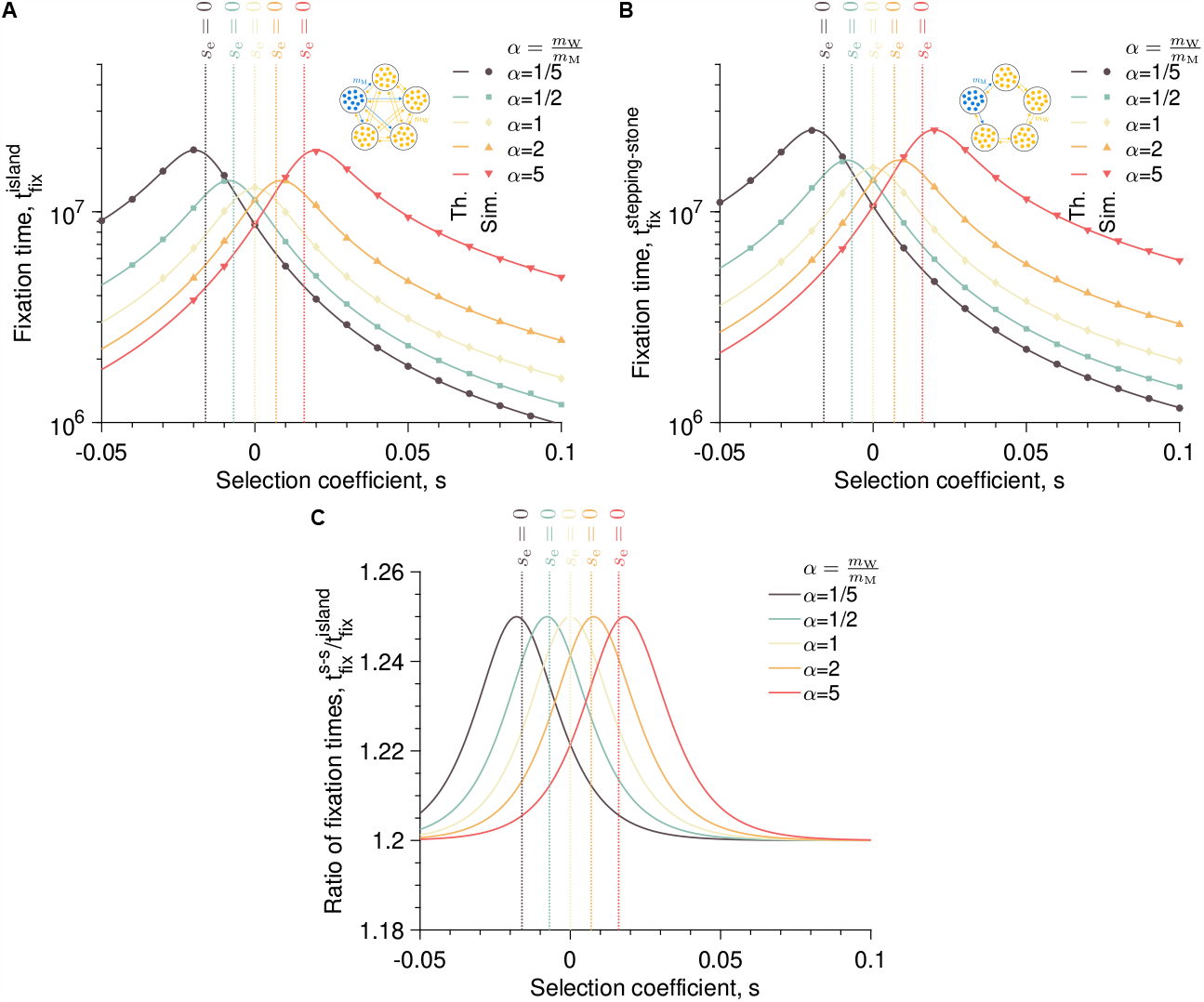
The fixation time allows for distinguishing the island and the stepping stone models. **A** The fixation time in the island model 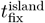 starting from one fully mutant deme is plotted against the selection coefficient *s* for different ratios *α* = *m*_W_*/m*_M_ of the wildtype and mutant migration rates. **B** The fixation time 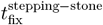 starting from one fully mutant deme is plotted against the selection coefficient *s* for different ratios *α* = *m*_W_*/m*_M_ of the wild-type and mutant migration rates. **C** The ratio of the fixation times 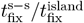 (with s-s standing for stepping-stone) starting from one fully mutant deme is plotted against the selection coefficient *s* for different ratios *α* = *m*_W_*/m*_M_ of the wild-type and mutant migration rates. In all the panels, the solid lines represent the analytical predictions (“Th.” standing for “Theory”), the markers are simulated data averaged over 10^4^ replicates (“Sim.” standing for “Simulations”), and the vertical dotted lines show the effectively neutral case (i.e., *s*_e_ = 0, where *s*_e_ = *s* − log(*α*)*/N* with *α* = *m*_W_*/m*_M_). Parameter values: number of demes *D* = 5, deme size *N* = 100, wildtype birth rate *b*_W_ = 1, death rate *d* = 0.1, wild-type and mutant migration rates for the island model (*m*_W_, *m*_M_) × 10^*−*8^ = {(2.06, 10.3), (4.11, 8.23), (6.17, 6.17), (8.23, 4.11), (10.3, 2.06)} (from top to bottom), wild-type and mutant migration rates for the stepping stone model (*m*_W_, *m*_M_) × 10^*−*8^ = {(4.12, 20.6), (8.23, 16.5), (12.3, 12.3), (16.5, 8.23), (20.6, 4.12)} (from top to bottom).

## Discussion

Whether on a microscopic or macroscopic scale, many populations are spatially structured. One type of spatial structure is a population subdivided into sub-populations, between which individuals can migrate, a process akin to gene flow. Whereas we understand the evolutionary dynamics of well-mixed populations, those of spatially structured populations need to be better understood. In this work, I quantified the impact of genotype-dependent gene flow on the fixation dynamics of a mutation in a population structured as the island or the stepping stone model. In particular, I combined analytical and numerical tools to compute the fixation probability of a mutation, the number of (fixed) migrants until the fixation of a mutation, and the fixation time of a mutation.

### The fixation probability of a mutation in the island and the stepping stone models exhibits an effective population size and selection coefficient

While evolutionary dynamics in subdivided populations have received much attention, much theoretical work has focused on the impact of population structure topology and gene flow pattern [14, 20]. One of the original features of my work is that I have introduced genotype-dependent gene flow by making the wild type and mutant have different migration rates. Deriving an equation for the fixation probability of a mutation in the island and stepping stone models and comparing it to the one in a well-mixed population allowed me to identify an effective selection coefficient and population size. Specifically, the effective selection coefficient differs from the selection coefficient if the wild-type and the mutant migration rates are different, whereas the effective population size equals the total census size. The effective selection coefficient shows that it is essential to consider the ability of a genotype to migrate when assessing its fitness. As a matter of example, a genotype reproducing less than another may still have greater fitness if it migrates more often and, thus, may take over the meta-population. This result confirms that, in some cases, gene flow can favor the fixation of locally deleterious mutations and, thus, limit natural selection [4].

For a long time, one of the key quantities for assessing the impact of a spatial structure on the evolutionary dynamics of a subdivided population has been the effective population size [15, 16]. Specifically, comparing the effective population size to the total census size allows for classifying the corresponding spatial structure as increasing or decreasing the efficacy of natural selection (i.e., a structure increasing the fixation probability of beneficial mutations and decreasing that of deleterious mutations or the other way round, respectively) [17, 18]. However, whereas it is well established that spatial structure can make the effective population size larger or lower than the total census size, its impact on the effective selection coefficient has been less studied. Yet, a theoretical work in which a diffusion approximation was applied to an island model with many demes exchanging migrants found that the population structure increases the effective population size and reduces the effective selection coefficient while keeping their product equal to the product of the total census size and the selection coefficient (i.e., *N*_e_*s*_e_ = *N*_tot_*s*, but *N*_e_ *> N*_tot_ and *s*_e_ *< s*) [19]. This result explains why the fixation probability in a subdivided population is the same as in a well-mixed population under some conditions [8, 9, 10, 11, 12]. This result also underlines that considering the effective population size alone is not enough to understand the impact of a spatial structure on the fate of a mutation but also requires the effective selection coefficient. In my study, I found that the effective population size is equal to the total census size in the island and stepping stone models. In other words, the spatial structure of these models does not increase or decrease the efficacy of natural selection. In contrast, the spatial structure can increase or decrease the effective selection coefficient, making the product *N*_e_*s*_e_ either larger or lower than *N*_tot_*s*, thus impacting the fixation probability of a mutation.

### Genotype-dependent gene flow increases the number of migrants until mutation fixation and the fixation time

The fixation time is often more difficult to calculate analytically than the fixation probability (see, e.g., [11, 17]). Yet, the fixation time allows for assessing the timescales involved in evolutionary dynamics, which can be crucial when evaluating the adaptation ability of a population undergoing environmental changes. In this work, I calculated the number of migrants until mutation fixation and the fixation time, given that the mutation becomes fixed. Interestingly, I found that genotype-dependent gene flow can increase the number of migrants and the fixation time.

In a well-mixed population, the fixation time of a beneficial mutation is the same as for a deleterious mutation with the same strength of selection (i.e., |*s*|) [9], although a deleterious mutation is very unlikely to fix. My model allowed me to extend this symmetry to subdivided populations with genotype-dependent gene flow such that the fixation time of beneficial and deleterious mutations are equal if they have the same effective strength of selection (i.e., *s*_e_). Also, whereas the population structure impacts the fixation time in the same way as the effective population size (e.g., a decrease in the effective population size decreases the fixation time) [17], I showed that genotype-dependent gene flow induces an increase in the fixation time. More specifically, the larger the difference between the wild-type and the mutant migration rates, the longer the fixation time.

### The fixation time allows for distinguishing the island and the stepping stone models as opposed to the fixation probability

My study and others showed that the fixation probability does not always allow for distinguishing different population structures. For example, the island and the stepping stone models give the same fixation probability, which makes them indistinguishable [14]. In this work, I derived an expression for the fixation time of a mutation in the island and the stepping stone models under the low migration limit. I showed that both models have different fixation times. More specifically, the fixation time is longer in the stepping stone model than in the island model, which can be explained by the number of migration paths [39].

This result was already numerically established in evolutionary graph theory [40]. This theory involves discrete-time models in which a subdivided population is structured on a graph with one individual at each node and probabilities that their offspring replaces a neighbor along each edge. The fact that this theory involves discrete-time models implies choosing dynamics, i.e., whether the first individual selected at each iteration reproduces or dies [41]. The problem with this theory is that the choice of dynamics impacts the evolutionary outcome [42], which raises universality issues. My model overcomes this choice of dynamics by considering continuous time, uncoupled reproduction, death, and migration events, in addition to considering sub-populations of variable size instead of single individuals. Moreover, my model allows for setting the total migration rate, enabling a rigorous comparison between different spatial population structures. Also, I went beyond numerical resolutions by deriving analytical predictions for the fixation time in the island and the stepping stone model (see [43] for numerical calculations in evolutionary graph theory).

### Theoretical perspectives

My work opens the door to several extensions. A first extension would be to consider other spatial structures and quantify the impact of genotype-dependent gene flow on the effective population size and selection coefficient. In the case of a continent-island model, in which a central deme is connected to peripheral demes, the effective population size could differ from the total census size if migration is asymmetric [14], but it is difficult to predict what value would take the effective selection coefficient. A second extension would be to relax the low migration limit hypothesis and investigate to what extent the predictions made in this work hold in the intermediate and strong migration limits. A third extension would be to adapt my model to make the link with experimental data more obvious. For example, in [44], the authors investigated the impact of asymmetric migrations on the fixation of a mutation in a spatially structured population by building a model inspired by batch culture experiments.

### Experimental perspectives

I believe my work can open the way to more quantitative comparisons between theoretical predictions and experimental results on the evolutionary dynamics of subdivided populations. Although experimental studies have focused on the impact of population subdivision on the magnitude of adaptation change [45, 46, 47, 48, 49, 50, 51], and the emergence and persistence of diversity [52, 53, 54, 55], more recent work has investigated its impact on the fixation probability and time [56]. Additional experiments could be performed with microfluidic devices [57], which would regulate gene flow between different sub-populations, or with microtiter plates [49, 50, 55], allowing migrations to be controlled with a liquid-handling robot.

## Acknowledgments

LM thanks the THEE Group and Kimberly Gilbert at UniBe for discussion and feedback on the manuscript. LM is grateful for funding from ERC Starting grant no. 804569 (FIT2GO) and SNSF Project grant no. 315230_204838/1 (MiCo4Sys) allocated to Claudia Bank (PI of the THEE group).

